# Modularity in the evolution of visual signals associated with aggressive displays

**DOI:** 10.1101/2024.02.22.581650

**Authors:** Kristina Fialko, Trevor D. Price

**Author notes:** **Corresponding author:** Address: Dept. of Ecology & Evolution, 1101 East 57th Street, Chicago, IL 60637.

## Abstract

Interactions between conspecifics commonly involve the use of stereotyped display movements, which can vary markedly between species. Theoretically, sexual selection by female choice can lead to large differences between species, but sexual selection by male competition may result in more limited diversification. Here, we evaluate display evolution in the aggressive signals of 10 leaf warbler species. Using high-speed videography of territorial behavior, we quantify differences in wing motion intensity and form. We find that both the rate of wing motion and the form of the display remain similar across species, which we attribute to an effective signal maintained through multiple speciation events. Differences among species arise though discrete additions to the behavioral repertoire (three species), loss of display (one species) and the presence of a pale patch on the wing. While some habitats differ discretely and dramatically in light intensity, this cannot account for all the differences in display behavior. We conclude that display evolution proceeds largely in a modular fashion. The basic conventional signal is maintained across species, enabling modifications to appear without loss of efficacy.

## INTRODUCTION

Displays to conspecifics are employed in a wide range of social situations including, during the breeding season, to attract mates (Mitoyen *et al*. 2019) and repel competitors (van Staaden *et al*. 2011). Spectacular displays are associated with sexual selection by female choice, and such displays often differ dramatically among closely related species. In birds, this is exemplified by the striking secondary sexual traits and associated displays of males in polygynous taxa, such as the Birds of Paradise (Scholes 2008, Scholes *et al*. 2017, Ligon *et al*. 2018, Miles & Fuxjager 2018), manakins (Prum 1990, Anciães & Prum 2008), and hummingbirds (Clark *et al*. 2018, Simpson & McGraw 2019). Models of diversification in such cases include co-evolution of male and female trait following slight displacements from equilibrium (Lande 1981, Kirkpatrick 1982) and mutation-order selection, whereby different attractive mutations arise in different populations (Price 2002, Mendelson *et al*. 2014). The implication from both the bewildering diversity of secondary traits in such groups, and from these models, is that display behaviors may shift in arbitrary, unpredictable directions, with little connection to differences among environments (Fig. 1A).

**Figure 1:**
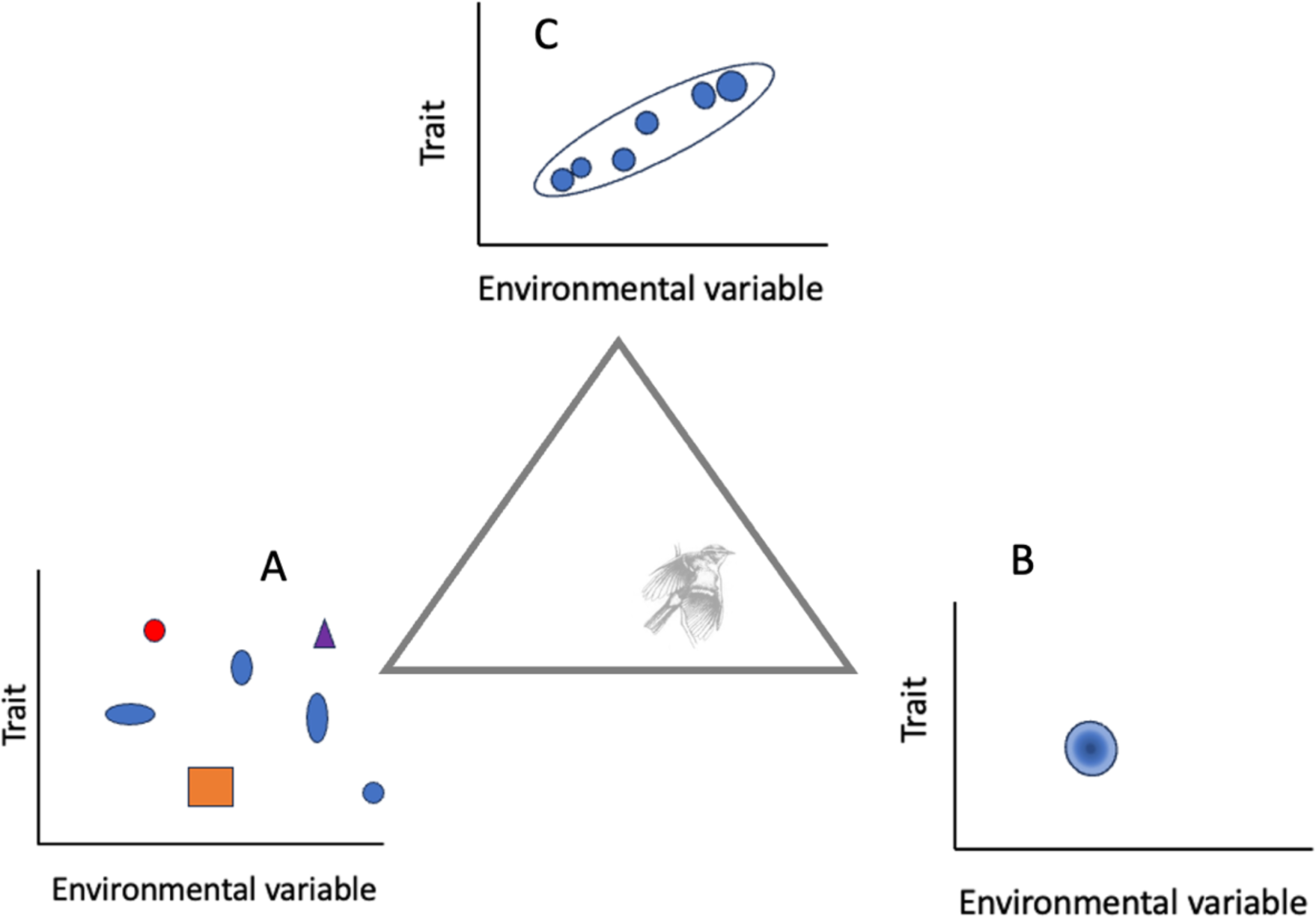
We consider three possible processes affecting the evolution of sexually selected traits. Anticlockwise from bottom left. (A) A collection of quite different displays accumulates among species, exemplified by sexual selection through female choice in polygynous species, with little influence of environmental differences. (B) Because traits used in competition at short range are expected to be optimized to be unambiguous and striking, all species inherit a similar display through their common ancestor. (C) The form of the display is modified according to environmental conditions (e.g. through sensory drive). The drawing indicates our findings. First, we show a basic display is conserved across species, as in (A). Second, we show some species have added unique components to this basic display, including some which can be related to environment (plumage patch) and others (a second display or loss of display), whose origins are less clear.

In contrast to displays used to attract mates, we might expect aggressive displays between males to be simple to transmit an unambiguous signal of intent (Morris 1957, Cullen 1966, Hurd & Enquist 2001). Further, such displays should often be short, so that individuals can display sequentially, and thereby assess each other (Catchpole 1980). While male competition can drive evolution through a number of mechanisms (Tinghitella *et al*. 2018), short simple displays may restrict possibilities for divergence. Instead, if the display is optimized to efficiently communicate between conspecifics, it may be passed little changed through descendants, and exhibit substantial stasis (Fig. 1B). For example, in *Anolis* lizards, territorial displays directed to males are highly stereotyped within species and differ in relatively small ways between species (Ord and Martins 2006).

Spectacular divergence in arbitrary directions and extreme stasis lie at ends of a continuum. Modifying both processes are effects of the environment (Fig. 1C). For example, despite large differences among related Birds of Paradise in the form of their displays, species found on the forest floor have larger display repertoires than those in the canopy (Ligon *et al*. 2018, Miles & Fuxjager 2018). And despite the small differences among *Anolis* lizards Ord and Martins (2006) were able to rank those differences (e.g., in dewlap pulse rate) to show how different features of the display correlate with shade vs. sunny habitats, number of sympatric species, occupancy of the canopy, and sexual size dimorphism. This and other studies of intrasexual aggression (Jenssen 1977, Fleishman 1992, Clark *et al*. 2015) suggest that although diversification may be quite limited, a relatively high fraction of the diversity may be a result of adaptation to different environments, associated with selection for efficient communication. Consequently, the evolution and adaptive significance of aggressive displays may be investigated using the comparative method, which seeks correlations between display and a species morphology and ecology (Fig. 1C).

The best known hypothesis that relates environmental differences to features of display is that of sensory drive (Endler 1992, Cummings *et al*. 2018). In this hypothesis, environmental differences impact transmission and perception, resulting in the evolution of traits involved in communication. Sensory drive has been most often applied to the evolution of color and color patterns, with one of the clearest examples being that of plumage brightness among *Phylloscopus* warblers breeding along an elevational gradient in the west Himalaya. Marchetti (1993) showed the brightness of an unpigmented patch on the wing (termed the wing-bar) correlates positively with darkness of habitat, which she inferred to result from evolutionary adjustments to maintain a certain level of conspicuousness. *Phylloscopus* males compete for territories (Marchetti 1998, Scordato 2018), utilizing short-range aggressive displays composed of rapid wing movements, which expose the wing bar (Marchetti 1993). Rapid wing movements are one of the most commonly used motions during avian threats (Andrew 1956, 1961, 2008, Tinbergen 1960, Kenyon & Martin 2022), presumably because the wings are so easily moved, with altered rates and overall form of wing motion potentially indicating aggressive motivation, increasing conspicuousness, or a combination of factors. We set out to ask if sensory drive has affected evolution of wing movements in the *Phylloscopus*, as brightness is expected to impact perception of motion-based signals as well as colors (Warrant 1999, Boström *et al*. 2016).

We address the following questions:

1. Is there substantial conservation of display form (Fig. 1B)?
2. Have discrete elements of display been added or lost across species (Fig. 1A)?
3. Is there any evidence for diversification in display associated with habitat (Fig. 1C)? For example, broader, more exaggerated wing movements may be present in darker habitats to enhance visibility.

We find evidence in support of stasis of the main display, punctuated by unique evolutionary events that result in the addition of a novel display or complete display loss. The presence of discretely different behaviors which have evolved just once precludes the strong use of the comparative method and suggests that a common mode of display evolution is modular, with addition of unique elements. We draw on environmental differences between species, plus previous work on the color patterns of these species (Marchetti 1993), to make some adaptive hypotheses about display evolution.

## MATERIALS AND METHODS

### STUDY SYSTEM

The genus *Phylloscopus* (leaf warblers) contains 76 species (Alstrom *et al*. 2018), which vary in mass from 5-12 g. In all species, individuals spend much of their time foraging for insects in trees and bushes (e.g., in a non-breeding season study, *P. trochiloides,* consumed one arthropod every 14 seconds throughout the day (Price 1981)). We studied 10 species that breed along a limited elevational gradient (2,000-4,000m; Price *et al*. 1997; Fig. 2) in the west Indian Himalayan state of Himachal Pradesh (Price *et al*. 2003). During the breeding season, species partition themselves along this elevational gradient in association with dominant plant species (Price 1991), resulting in occupancy of distinct primary breeding habitats (Fig. 2). All species are partial or complete migrants, spending the non-breeding season at lower elevations and latitudes.

**Figure 2.**
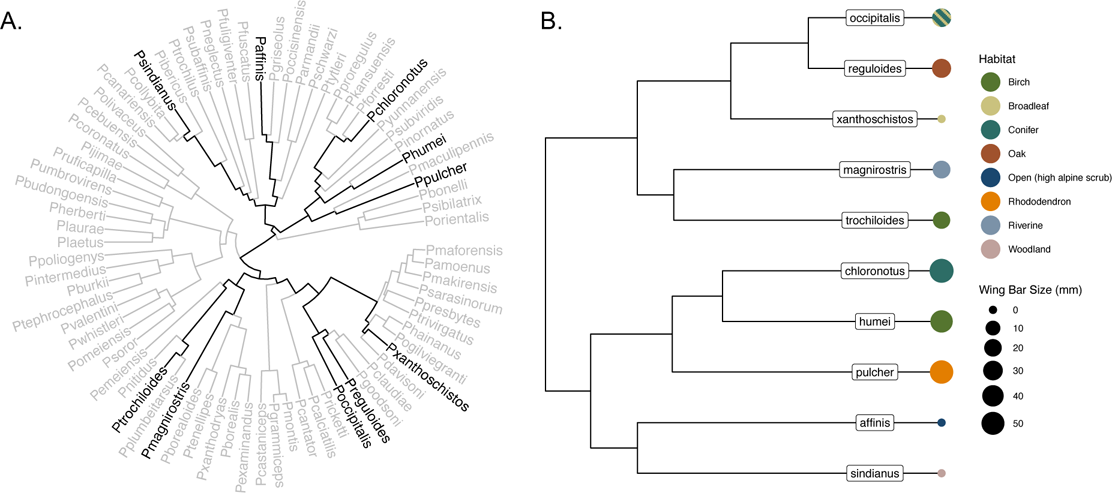
*Left Phylloscopus* phylogeny (Alstrom *et al*. 2018), with the 10 species in this study highlighted in black. Note that the species we study span the root of the tree. *Right.* Size of the point at the tip corresponds to each species wing bar size (Price & Pavelka 1996) and is color coded by primary habitat during the breeding season; note the two species pairs which show convergent evolution in habitat. Constructed using ggtree (Yu *et al*. 2017)

All species have similar plumages, possessing greenish-olive to brown upperparts and pale underparts. Many species have a light stripe of unmelanized feather keratin across the tip of the greater covert feathers, producing a wing bar. Wing bar size varies between species (Fig. 2, (Price & Pavelka 1996) and for the one species where it has been studied (*P. humei)* wing-bar size is about 10% larger in males than females (Scordato *et al*. 2012). Color of the sexes is similar, as assessed spectrophotometrically (*unpublished data).* Plumages do not vary seasonally, except for feather wear, which can reduce the size of wing bars over time (Scordato *et al*. 2012).

We conducted simulated territorial intrusions using playback experiments and filmed species responses in the breeding season. To quantify variation in wing motion, we applied a geometric morphometric approach to compare wing trajectory shapes within a morphometric space. To assess the influence of the light environment we measured habitat illuminance in the primary breeding habitats across the elevational gradient.

### FIELD METHODS

Author 1 studied warbler behaviors during the breeding season in the Manali Wildlife Sanctuary, Himachal Pradesh, India (32.25°N, 77.17°E, spanning 2000m - 3600m, between April 22 –June 22, 2019 and April 25 – May 30, 2022) and at Nain Gahar village, Himachal Pradesh, India (32.73 °N, 76.86 °E between June 11-21, 2019 and June 15 – July 8, 2022). They also visited two sites in Arunachal Pradesh (26.97 °N, 92.92 °E and 27.06 °N, 93.03 °E) and one site in Andhra Pradesh (17.81 °N, 82.49 °E) during the non-breeding season (December 22, 2021 – January 22, 2022). We collected two sets of data: the first on rate of wing movement during foraging and territorial intrusions, and the second on form of the display (Table 1). Rate of wing movement was collected because it was qualitatively apparent that movement increases during aggressive responses (call note rate similarly increases, (Wheatcroft 2015) and because variation in display rates may indicate differences in aggressive motivation and condition (Yasukawa 1978, Clutton-Brock & Albon 1979, Ord & Evans 2003, Barnett *et al*. 2014). Display form was collected to quantitatively assess whether displays vary using a geometric morphometric approach and to explicitly address the question of how displays have been modified across species.

**Table 1:** Sample sizes from the video data. Foraging videos (column 2) were taken opportunistically through the breeding season (April – July, labeled with *b*) and a few from the nonbreeding winter season (December – January, labeled with *w*). Behavioral trials were filmed using a single 60 fps camera (column 4) and a 480 fps high-speed camera array (column 5) and were all conducted during the breeding season. Many individuals readily performed territorial wing displays (column 3) but only a subset of these were captured on video used in data analysis. Double, single and shiver wing flicks in column 6 refer to three qualitatively different displays, as described further in the results section.

| Species | Foraging videos (n) | Behavioral trials with aggressively responding males (n) | Territorial 60 fps video recordings used in rate analysis (n) | Territorial high-speed (480 fps) video recordings (n) | High-speed videos with a lateral orientation used in shape analysis |
| --- | --- | --- | --- | --- | --- |
| *P. affinis* | 3 (2 *b*, 1 *w*) | 11 | 5 | 5 | NA |
| *P. chloronotus* | 3 *b* | 16 | 5 | 11 | 7 double |
| *P. humei* | 4 *b* | 17 | 4 | 13 | 5 double |
| *P. magnirostris* | 0 | 5 | 0 | 3 | 3 double |
| *P. occipitalis* | 3 *b* | 15 | 2 | 10 | 3 double, 3 single |
| *P. pulcher* | 4 *b* | 18 | 2 | 12 | 2 double, 7 shiver |
| *P. reguloides* | 5 (3 *b*, 2 *w*) | 12 | 2 | 7 | 2 double, 1 single |
| *P. sindianus* | 2 *b* | 4 | 2 | 3 | 1 double |
| *P. trochiloides* | 5 (1 *b*, 4 *w*) | 6 | 2 | 2 | 2 double |
| *P. xanthoschistos* | 4 *b* | 11 | 3 | 6 | 3 double |

To document the use of wing movements during foraging, we opportunistically filmed individuals. When a species of interest was detected, we used a single Sony RX10 DSC III camera mounted on a tripod set at 60 frames per second (fps) and filmed the individual for as long as possible. Birds were identified to species at the time of filming or when reviewing the video footage through call notes, songs, plumage, or a combination of these traits.

Most of the data comes from territorial playback experiments in the breeding season. Author 1 located singing males between 0500-1100. Experiments consisted of two parts. During Part I we established a filming focal point and observed the territory owner’s response. We placed a Bluetooth speaker (Ultimate Ears WONDERBOOM) in the target individual’s territory and played that species song to simulate a territorial intrusion. If the territory owner responded by singing back and approaching the speaker, we continued to play the song for 5 minutes to observe where the individual would perch in the territory. This was to maximize the likelihood that the camera setup would capture the behaviors of interest. In total, we attempted 306 behavioral trials. Of those, 43% (n = 132) were terminated during Part I due to either poor filming conditions or lack of response from the territory owner.

Part II: Once a consistent focal point was established, we set up two camera teams. The first team consisted of a single person with a camera (Sony RX10 DSC III) mounted on a tripod, filming the target bird at 60 fps continuously during the trial. This allows for an extended view of the display and was used to calculate wing flick rate and record all the motions present in the species display repertoire. The second camera team operated a high-speed camera assembly, which consisted of three Sony RX10 DSC III cameras mounted on tripods, each equipped with a Ziv TRS-10 Timer Remote set to the same channel, allowing for simultaneous remote triggering. We used three cameras to increase the chances of capturing displays during which the bird is oriented laterally (defined as the line from beak to tail running perpendicular to the camera lens). The cameras were placed at least 3 m. from the focal point, in an arc with each camera separated by 45° from the next one. The cameras were set to film at 480 fps on a delayed trigger. This mode continuously films until the trigger is pressed, at which point the prior two seconds of footage are written to the SD card, thereby allowing the capture of display behaviors without a response delay from the observer. Both setups were left undisturbed for 10 minutes before the trial started. During the trial the two observers remained 8 meters away from the focal point.

During the behavioral trials, Author 1 played a target species song for 10 minutes. In 34% of trials (n = 174 total trials) the bird did not respond, at which point the trial was deemed unsuccessful and ended (n = 59 terminated trials). For clarification, this differs from the termination described in part I; here the individual responded during the pre-trial period (Part I) but then ceased responding after the camera array was set up and trials began. Because of set-up time and the 10-minute undisturbed period, approximately 20-25 minutes could elapse between song playbacks.

If the bird responded by calling or singing back or by approaching the speaker during this time, the trial continued. The high-speed camera assembly was triggered by Author 1 when the bird perched near the focal point, within the camera’s frame of view. The single camera team filmed the bird for as long as possible until sight of the bird was lost, at which point camera recording was paused. The playback trial continued until the end of the 10-minute period, accumulating as many 2 second videos as possible when the bird displays at the focal point. If the bird continuously displayed throughout the time of the first trial, we would leave the camera array in place and begin a 5-minute pause period. After the pause period, the trial was repeated up to a maximum of three times. In total, we had 115 aggressively responding males distributed across 10 species and were able to record high speed (480 fps) videos of 73 males and 60 fps videos of 68 males (Table 1).

After the behavioral trials, we calibrated the filming area. We placed a 3’’x 5’’ checkerboard and XRite ColorChecker Passport close to the area where the bird perched. We then moved the checkerboard through the filming area, pointing to each of the camera views. These tools were used to provide a scale visible from any camera angle.

### LIGHT MEASUREMENTS

To assess the influence of the light environment on wing displays, we measured habitat illuminance in the primary breeding habitats across the elevational gradient. In both breeding season field sites (Manali, Himachal Pradesh and Nain Gahar, Himachal Pradesh), we deployed Onset light and temperature loggers (HOBO Pendant MX2202 Temperature/Light Data Logger), resulting in 5-13 samples per habitat across the two field locations (Table S1). Light loggers were mounted horizontally with the light sensor facing the sky on either horizontal branches in woodlands (birch, conifer, rhododendron, oak) or on PVC pipes staked in the ground in understory and open habitats. The placement of loggers reflected where the associated species were commonly found; canopy or midstory in wooded areas and low-lying shrubs or the ground in open or understory areas. Loggers were configured to record light and temperature every minute. Data from the loggers were downloaded to the HOBOconnect app via Bluetooth at the time of collection.

## ANALYSIS

### WING MOTION

We quantified variation in wing motion using two measurements: rate (from the 60 fps videos) and form (from the 480 fps videos). From the single 60 fps camera, we measured wing flick rate in two contexts: foraging and territorial response. Videos were selected for foraging analysis if the individual could be seen actively searching for or capturing prey. Territorial responses were filmed at the time of simulated territorial intrusion experiments. We calculated rate as the number of wing flicks an individual performs divided by the total amount of time the bird is present on screen. Using the trim function on QuickTime Player (Apple Computer) with video playback at half speed, we analyzed the video frame by frame to quantify wing flicks. To separate flicks from wing movements used in locomotion we only counted wing flicks when the bird hopped less than one body length during the observation sequence. Total display time is the difference in the timestamp when the individual leaves the frame of view and the timestamp when the focal individual first appears in the frame. We only selected individuals that completed at least 5 wing flicks during the time on video. For individuals that left the frame and reappeared over the course of a video recording we took the first display sequence in which the bird performed at least 5 wing flicks.

To quantify the form of the display during the territorial playback experiments, we used the high frame rate videos in which the bird is in a lateral orientation. We determined lateral orientation visually and then confirmed orientation by viewing the angle from the other two cameras. This resulted in a sample size of high-quality recordings from 37 individuals from 10 species, reduced from a total high-frame rate dataset of 73 individuals. We extracted frames using the extractFrames function in StereoMorph (Olsen & Westneat 2015). We isolated each individual wing flick, defined by the frame in which the wing begins the upstroke to when it returns to the starting position after the downstroke. We numbered the frames from 1 (initiation) to the end of the flick; the number of frames varied from 35-45. To compare the variation in the form of the display while accounting for variation in duration, we described the shape of each wing flick using 15 time points (Fig. 3). First, we identified three specific wing positions from each video: initiation, end and maximum wing extension. The initiation and end of the wing flick were assigned time points 1 and 15, respectively. The point of maximum wing extension, when the wing transitions from the upstroke to the downstroke was assigned time point 7 (Fig. 3). From these we added 12 additional time points by extracting frames uniformly dispersed between 1-7 (5 points) and 8-15 (7 points).

**Figure 3.**
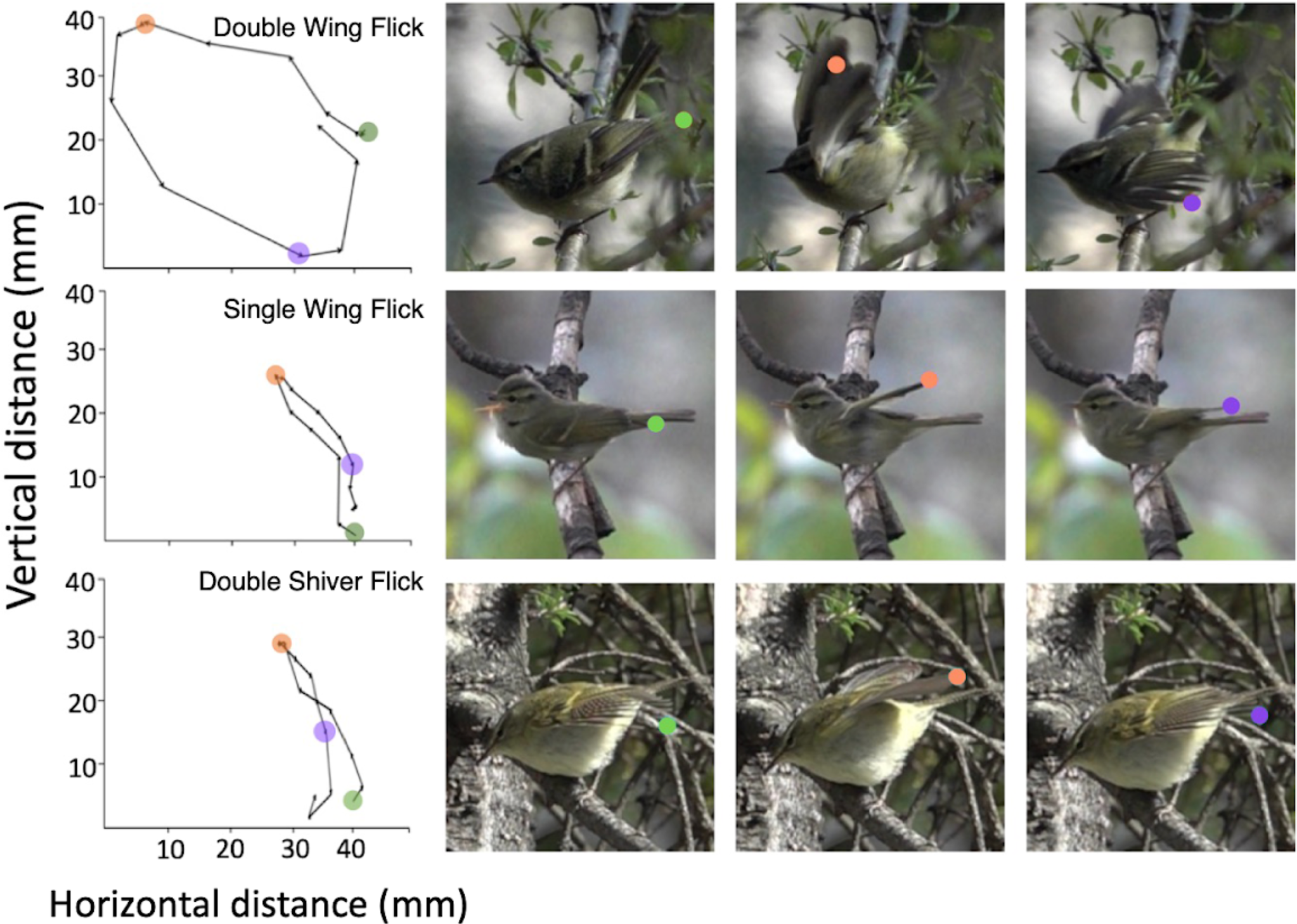
*Left* Trajectories for the displays observed in the warblers studied. Plotted is the distance moved in 15 equal time intervals. The dots correspond to 3 time points. L1 (green): when the wing flick begins. L5 (orange): when the wing reaches the upstroke:downstroke transition. L13 (purple): A sample landmark showing how this position can differ depending on the type of motion used in display. *Right* Video frames for each labeled point. From the top, species are *Phylloscopus chloronotus*, *P. reguloides*, *P. pulcher.* Video examples of these behaviors can be found in the supplementary data.

All statistical analyses were conducted in R, version 4.2.2 (R Core Team 2022). We used the labelFrames function in StereoMorph to place a landmark on the tip of the 8th primary feather for each of the 15 frames. Each wing flick is then described using 15 “homologous” landmarks, creating a shape capturing the trajectory of wing motion (Fig. 3). The landmarks describing the trajectory shapes were scaled and aligned using the Generalized Procrustes Analysis (GPA) in the R package borealis (Angelini 2022) to remove variables of size, rotation and orientation, leaving a set of aligned coordinates that capture variation in shape. We performed a principal component analysis (PCA) on the correlation matrix of the aligned coordinates and visualized the location of each individual’s trajectory by plotting the first two PCA axes. We plotted convex hulls around the data points in the morphospace generated by PC1 and PC2, the most significant axes of variation. We used lme4 (Bates *et al*. 2015) to calculate percent variance between and within species display components for both PC1 and PC2.

To visualize how the shape of wing movement changes along PC1 and PC2, we back transformed the PCscores to their relative positions in the morphospace (Olsen 2017). The points along the outside of each backtransformed shape correspond to the 15 landmarks used to describe the trajectory.

### LIGHT MEASUREMENTS

We extracted the lux values from the logger files and log-transformed the data. To standardize for longer day lengths as the season progressed, we filtered the data to include times between 0600 -1800, which is a time interval occurring after sunrise and before sunset through the entirely of the breeding season. We then took the average lux measurement per logger per day. Results from habitats present in both locations were similar and we combined them. We fitted a linear mixed-effects model with habitat as a fixed effect and location and logger ID as nested random effects using lme4. We then performed a post-hoc pairwise comparison of the habitats using the R package emmeans (Lenth 2023).

## RESULTS

We observed three discrete behaviors (Fig. 3). One behavior, which we term the *double wing flick* is shared between all species. This behavior resembles motions used during takeoff, where both wings are moved simultaneously through rotation at the elbow and shoulder joints during the upstroke. During the downstroke, the humerus is extended horizontally from the body, resulting in the extension of the distal portion of the wing until it folds back to rest near the starting position. One species *(P. pulcher)* commonly conducts a *double shiver flick,* distinguished from the double flick by both shape (Fig. 5B) and rate (Fig. S1). During this behavior the wings are extended horizontally from the body at the shoulder and undergo a series of rapid rotations at the elbow and wrist joint, resulting in a shivering motion. During territorial displays, this behavior is repeated and rarely interspersed with double wing flicks. Finally, two related species *(P. occipitalis* and *P. reguloides)* conduct a *single wing flick,* whereby one wing is raised vertically from the body at the elbow joint, but the wing does not undergo a horizontal extension. Instead, the wing is placed back to the starting position before alternating with the other. Both species regularly intersperse single wing flicks with double wing flicks during both foraging and territorial displays (Table S2).

**Figure 4:**
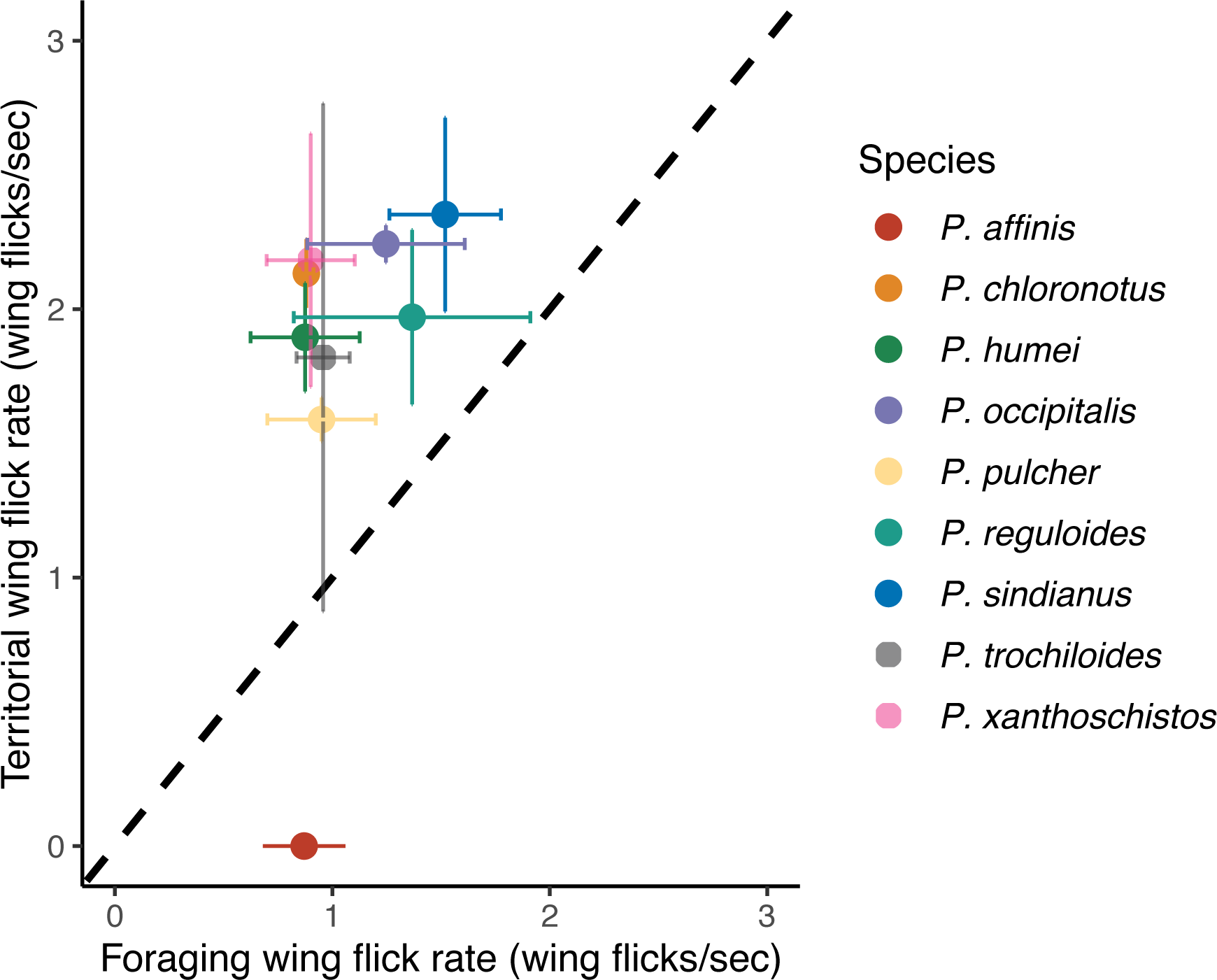
Species territorial wing flick rate plotted against foraging wing flick rate, with standard error. The black dashed line is the line of equality; 8 species flick wings faster in territorial interactions. Among the 9 species foraging and territorial rate are not correlated (r = 0.43, P = 0.2). For sample sizes for each species, see Table 1.

**Figure 5.**
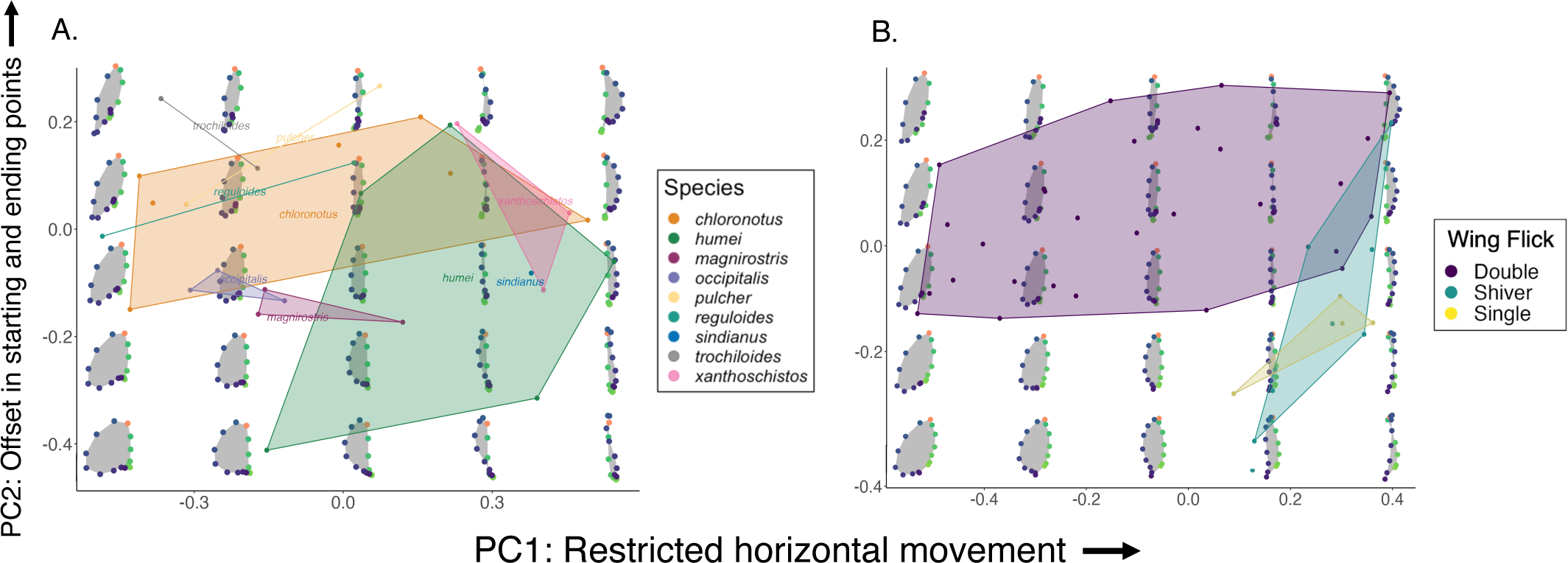
(A) Double wing flick shapes for individual males. The colored lines circumscribe the convex hull for each species (note that some species have a sample size of n = 2, so are connected by a line). Background (in grey) illustrates the trajectories; upstroke landmarks (points 1-6) are in green, the upstroke:downstroke transition (point 7) is shown in orange and downstroke landmarks (points 8-15) are in purple. PC1 (52% of the variance) describes a restriction in horizontal movement of the wing and PC2 (15%) represents an offset in the starting and ending points of the wing. (B) Principal components were conducted on the entire dataset for the three wing flick types. The convex hulls enclose all species for each wing flick type (double, as in the left plot, single: *reguloides, occipitalis*, shiver *pulcher*.) PC1 (46%) describes a restriction in horizontal movement of the wing and PC2 (13%) an offset in the starting and ending points of the wing.

### WING FLICK RATE

*P. affinis* does not flick its wings at all in response to aggressive playback, although it does so when foraging (Fig. 4). All other species flick their wings significantly faster during territorial displays than during foraging (Fig. 4, Table S3). All species, including *P. affinis,* have similar foraging wing flick rates (*F_8,24_* = 0.49, *P* = 0.8). Once *P. affinis* is excluded, species do not differ significantly in territorial display wing flick rate (*F_7,14_* = 0.44, *P* = 0.9, Table S4). Shiver flicks are the dominant behavior used by *P. pulcher* during territorial contexts, comprising 95% of the motions used during a display. Shiver wing flick rates are significantly faster than double wing flick rates (Fig. S1). Shiver flicks are only used during territorial interactions, and we observed no instances of this behavior during foraging. Within the foraging context, single wing flick rates do not differ significantly from double wing flick rates in *P. occipitalis* and *P. reguloides.* Single wing flicks used during territorial displays are significantly faster than those used during foraging (Fig. S2).

### WING TRAJECTORY

We first analyzed double wing flicks by the 9 species that use them during territorial displays. The primary axis of variation (PC1: 52% of the variance explained) describes reduced horizontal movement of the wing (Fig. 5A). Individuals with low values of PC1 move their wings more elliptically while those with high values of PC1 move it along a more constrained vertical axis. PC1 scores do not vary significantly among species (ANOVA: *F_8,19_* = 2.03, *P* = 0.10, table S6). The second axis of variation (PC2: 15% of the variance explained) corresponds to an offset of the starting and ending points. Individuals with low values of PC2 tend to place their wings close to the point at which they initiate their wing flick, while those with high values of PC2 have more variability with where the final downstroke points land relative to where they start. Species did not vary significantly in their PC2 scores (ANOVA: *F_8,19_* = 1.75, *P* = 0.15, Table S6). Most of the variance is within versus between species (75% within for PC1 scores, 79% for PC2 scores).

Next, we included all three wing flick types in a single analysis, where we combined observations from the 9 species to compare the three types. The three motions differ significantly along both PC1 (ANOVA: *F_2, 37_* = 9.24, *P* = 0.001, *a posteriori* tests are in Table S8) and PC2 (ANOVA: F_2,37_ = 7.98, P = 0.001, *a posteriori* tests are in Table S8). Shiver and single wing flicks have higher PC1 scores relative to double wing flicks (Fig. 5) because the upstroke and downstroke landmarks are closer together, with less horizontal motion. Both shiver and single wing flicks have lower PC2 scores than double wing flicks, reflecting more consistency in placing the wing tip back in same region it started. Although shiver flicks (n = 8) occupy a larger area of the morphospace along PC2 (Fig. 5B) than single wing flicks (n = 4) this may be due to a larger sample size capturing more individual variation.

### HABITAT LIGHT

Habitats differ in illuminance (Fig. 6); *a posteriori* pairwise tests indicate that *open* is brighter than all other habitat types and *understory* is significantly darker than all other habitat types (Table S10). Conifer, birch, oak, and rhododendron did not vary significantly in brightness between each other.

**Figure 6.**
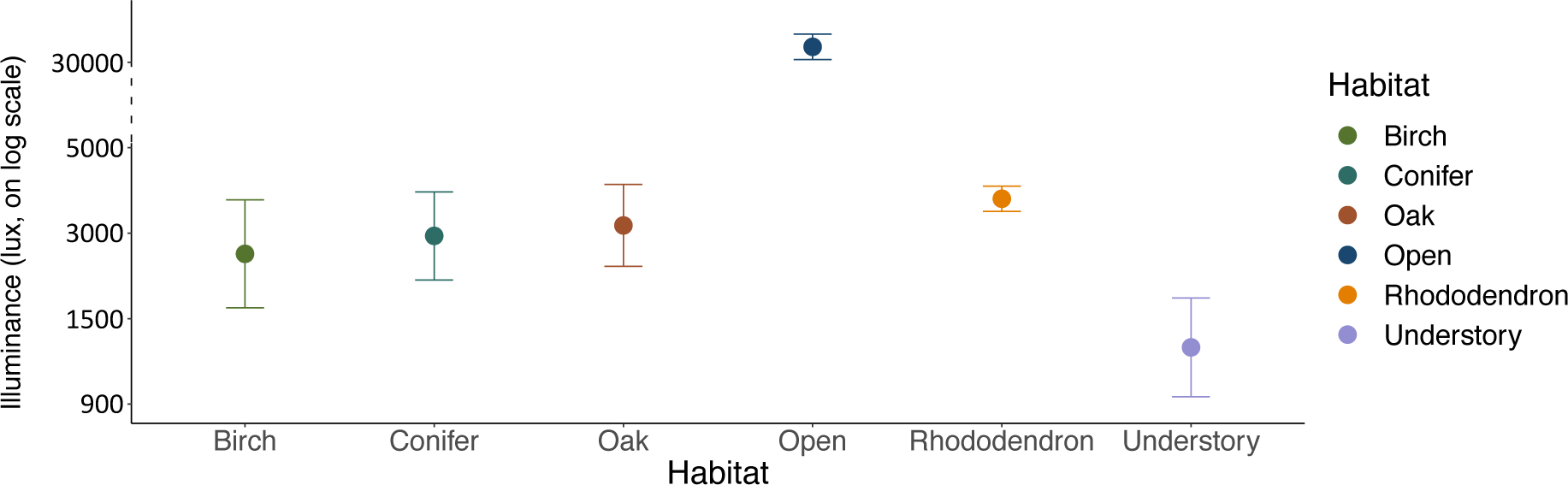
Brightness for 6 different habitats in the western Himalaya (mean ± standard deviation). For statistical tests see table S9. Pairwise Tukey tests indicate open and understory habitats are significantly brighter and darker, respectively, from birch, conifer, oak and rhododendron, which are not different from each other.

## DISCUSSION

Within species, individuals often communicate using specific and repetitive patterns of ritualized movements. In this study, we investigated the use of wing motion in the aggressive displays of *Phylloscopus* warblers to assess the extent of display variation and whether it is associated with habitat. We find that the primary display used in response to territorial playback, the double wing flick, remains conserved in both rate and overall form across species. Despite the widespread use of this display and its expected efficacy in close-range interactions, three species have modified their behavioral repertoires through the addition of novel, discrete behaviors: the shiver wing flick (*P. pulcher*) and the single wing flick (*P. occipitalis* and *P. reguloides*) and one species, *P. affinis,* has lost the display. This repertoire of wing flick behaviors align with descriptions in earlier studies (Marchetti 1993), and our analysis introduces a novel quantitative lens, offering a comparative perspective across species. These modifications are the result of three evolutionary events. Further, a pale wing-bar has been independently added twice during the divergence of these species from their common ancestor (Price and Pavelka 1996, Fig. 2), and is present in 7 of the species. We first evaluate why the primary display remains similar across species, and then investigate each of the modifications in turn. Finally, we evaluate the relationship between display and habitat.

### WING DISPLAY DIVERSIFICATION AND FUNCTION

Behaviors used in an aggressive signaling context may be under greater selective pressure to remain simple and consistent compared to behaviors used in courtship (Tinbergen 1960, Irwin 1996), which are shaped primarily by sexual selection through female choice Our results largely support this hypothesis, as the primary display used across species, the double wing flick, is found in 9 of 10 species in response to territorial playback. We applied a geometric morphometric approach to quantify variation in the shape of the wing display to test for subtle and continuous differences in display form. Instead, we largely found that most of the variation in the double wing flick is within species, not between, with 75% and 79% of the variation occurring among individuals within species than among species for PC1 and PC2 shape scores, respectively (Fig. 5). Species also do not vary significantly in their rate of wing flicking in territorial contexts (Fig. 4). This conservation in the double wing flick suggests that this behavior may be under little selective pressure to diversify.

Within a species, wing displays are variable across two measures – rate and form (Figs. 3 and 5). By comparing a signaling (aggression) and non-signaling (foraging) context, we showed that an aggressive stimuli (song playback) induces an increase in wing flicking rate, except for one species which drops the display altogether (Fig. 4). Observations of aggressive interactions between conspecifics confirm the use of a high wing flicking rate in territorial disputes, which, if unresolved, then escalate to chasing behaviors and can end in physical fights (Price 1981). An increase in rate compared to a nonsignaling context and an association with attack escalation imply the wing movements are likely used as threat displays (Számadó 2003). Display rate is associated with levels of aggressive motivation in many other taxa (Deag & Scott 1999, Lange & Leimar 2003, Ord & Evans 2003, Castro *et al*. 2006, Elwood *et al*. 2006, Brown *et al*. 2007, Crothers & Cummings 2015).

Potentially receivers may be assessing subtle information in form (Byers *et al*. 2010, Barske *et al*. 2011) but it is conceivable wing motion may have no information content but rather serve to amplify other traits, such as color patches (Hasson 1991, Bókony *et al*. 2006). Indeed, Marchetti (1993, 1998) showed the wing-bar of one *Phylloscopus* species *(P. humei)* functions in aggressive interactions. Motion is one of the most effective ways to capture attention (Abrams & Christ 2003, Franconeri & Simons 2003, Rushton *et al*. 2007) and in birds, wing movement is perhaps the simplest way to increase visibility. Further work is required to elucidate what specific features competitors are assessing during aggressive interactions, which should increase our understanding of why features of the display are evolutionarily constrained.

### ADDITIONAL DISPLAYS

In addition to the baseline double wing-flick, a subclade within the *Phylloscopus* have a display in which species alternate single wings (see Table S2 for example sequence) (del Hoyo *et al*. 2020a, b). The clade includes both *P. reguloides* and *P. occipitalis* from this study. The single wing flicks are often intermingled with double wing flicks during both foraging and aggressive displays (Table S2) and it is used by both males and females during foraging throughout the year. These species form large flocks in the winter (Macdonald & Henderson 2008, Hariharan *et al*. 2022), where wing motions may serve to facilitate flock cohesion or communication. The rate of the single wing flicks increases during male aggressive interactions (Fig. S2), so the display functionally operates in the same way as the double wing flick. However, the single wing flick is distinctive in its trajectory, where the distal portion of the wing remains unextended and most of the motion is concentrated in lifting the wing at the wrist joint. This results in relatively less horizontal movement than the double flick (Fig 5B).

The other display, employed by *P. pulcher* is different. This display, the shiver double flick, is used only in the breeding season and largely replaces that of the double wing flick although the double flick is still occasionally used during display bouts. The shiver flick is characterized by faster movement (Fig. S1), achieved through a series of rotations at the wrist joint (Fig. 3). This also results in relatively less horizontal motion than the double wing flick (Fig. 5B), although the motion itself is different from the single wing flick. It resembles the motions used by young birds of all species when they are begging for food (Supp. Video 1), but the reasons why it has been established as an aggressive display in this species alone remain obscure.

The addition of these discrete behavioral elements to aggressive displays rather than a replacement of the shared form mirrors results from aggressive contexts in other taxa. Comparative analyses of *Anolis* have documented the use of display modifiers, which are additional movements that are added to shared core displays (Jenssen 1977, Ord *et al*. 2002). Multiple forms of threat display may have evolved to reflect different escalatory steps (Andersson 1976, Hurd & Enquist 2001) or to overcome reduced reliability in the original signal (Andersson 1980). Alternatively, different display behaviors may mediate species recognition (Macedonia & Stamps 1994, Clark *et al*. 2015), as has been suggested for variation in other visual signals such as color (Couldridge & Alexander 2002, Klomp *et al*. 2017, Dyson *et al*. 2020). In our case, transmission of species identity seems unlikely as the displays are used at close range once a challenger has been identified as a conspecific. Further, the two closely related species with single and double wing flicks in their repertoire, are exceptionally similar in their plumage and morphology. Hence, they would be expected to have diverged in the display if it was evolved in species recognition.

### SENSORY DRIVE

In Kashmir, Marchetti (1993) observed that *Phylloscopus* species with wing-bars inhabited darker environments than those without wing-bars, which she attributed to sensory drive. She argued that in darker habitats, species maintain visibility by becoming brighter in appearance. *P. affinis* is the one species without wing-bars held in common between that study and ours. This species breeds above tree line in high alpine juniper (Price 1991), which has substantially higher illuminance than all other habitats we studied (Fig. 6), reflecting the open composition of this habitat with little to no tree cover. This species does not flick its wings in display (Marchetti 1993) suggesting a role for sensory drive in not only affecting plumage, but also display. The lack of wing flicking in this species is not a consequence of reduced aggressive responses. Indeed, in our experiment, the territory owner responded very aggressively, singing back, approaching, and even attacking the speaker, but it never flicks its wings. The signaling environment may provide some clues as to why it has dropped wing movements. A distinctive feature of *P. affinis* is the prominent yellow (carotenoid based) underparts (Grimmett *et al*. 2012 p. 340 plate 151). We suggest that detection and assessment in this species may be achieved by display of the underparts. It bears noting that two other species without wing-bars both flick their wings. One, *P. xanthoschistos*, has yellow underparts but lives in woodland, and the second, *P. sindianus* lives in relatively open habitats, whose light environment we were not able to measure but is likely to be intermediate between that of open habitat and dense woodland.

In dim light conditions the tradeoff between temporal and spatial resolution becomes exacerbated (Lythgoe 1979). We predicted that poor motion discrimination in dark environments can lead to pressures to exaggerate critical features of a motion-based display, which may lead to interspecific variation in the use of wing movement. However, with the exception of *P. affinis*’*s* habitat light intensity was similar across the habitat types occupied by other species. This differs from the results found in Kashmir by Marchetti (1993) and suggests that there may be geographic variation in habitat features across these species distributions. However, light intensity is only one axis upon which the sensory environment can vary. The spatial organization of the background (Hulse *et al*. 2020) and its motion (Ord *et al*. 2007, Peters 2013) are other variables that should affect the perception of a visual display and remain to be assessed in this system.

## CONCLUSIONS

Our ability to dissect display movements using high speed video coupled with a novel use of a morphometric approach to study bird displays shows that rather than diversify across species, the form of the primary aggressive display has been largely conserved. Aggressive displays are expected to be simple in form in order to convey an unambiguous message (Hurd & Enquist 2001), e.g. through changes in rate. Once such a display efficiently conveys a message it may be carried through subsequent speciation events. Nevertheless, we find that the primary display has been built on to generate differences among species, through either its complete loss, by the addition or subtraction of color patches on the wing, or by addition of qualitatively different displays. The effect of the signaling environment is weak, with the only possibility we highlight being the loss of display in one species. Given that environments vary in many ways other than light intensity, such as foliage structure and background color, we anticipate that future detailed studies of habitat will further our understanding of the origin of qualitatively different displays, and their link to color patch evolution. At present, however, we consider aggressive displays to have evolved through a mix of strong stabilizing selection on some elements, “arbitrary” addition of an effective display in some lineages, and mild influences of the habitat (Fig. 1).

