## Supplementary Data for "Modularity in the evolution of visual signals associated with aggressive displays"

**TABLE OF CONTENTS**

**SUPPLEMENTARY FIGURES	2**

Figure S1: Territorial double and shiver wing flick rates for *Phylloscopus pulcher.*	2

Figure S2: Single wing flick rates for foraging and territorial displays used by *P. occipitalis.*	3

**SUPPLEMENTARY TABLES	4**

Table S1: Sites and sample sizes for light loggers.	4

Table S2: Single wing flick sequences used by *P. occipitalis* and *P. reguloides.*	5

Table S3: Statistical test results for relationship between wing flicks per second, species, and context.	6

Table S4: Pairwise Tukey test results for species on territorial wing flick rate.	7

Table S5: PC1 and PC2 scores for double wing flick shapes.	8

Table S6: Pairwise Tukey tests for PC1 and PC2 scores of wing flick shape on wing flick type.	9

Table S7: Pairwise comparisons of the effects of habitat type on light (lux) from a linear mixed-effects model.	10

**SUPPLEMENTARY VIDEOS	11**

Description for supplementary video files	11

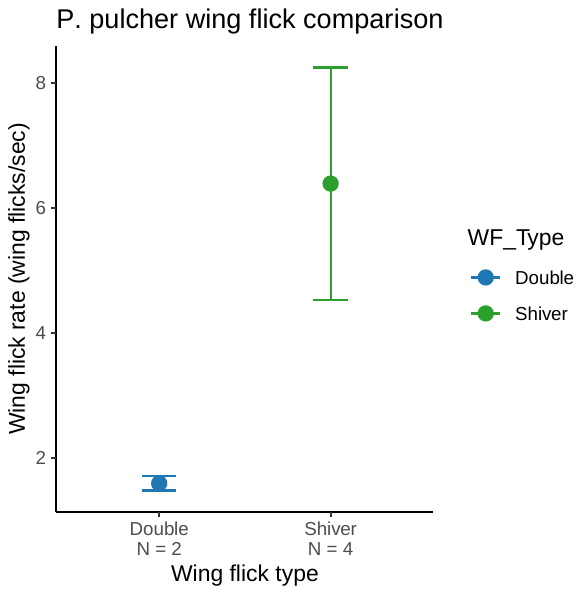

**Figure S1**. Territorial wing flick rates for the two types of wing flicks used by *Phylloscopus pulcher,* with standard deviations. Shiver flicks are faster than double flicks. (ANOVA: *F_1, 4_* = 11.82, *P* = 0.03).

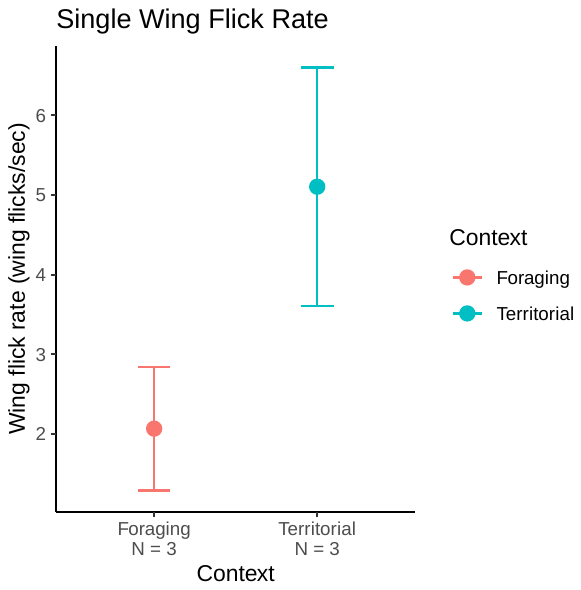

**Figure S2.** Single wing flick rates for foraging and territorial displays used by *P. occipitalis,* with standard deviations. Single wing flick rate is faster during territorial contexts compared to foraging (ANOVA: *F_1, 4_* = 9.719, *P* = 0.04).

| Site | Birch | Conifer | Oak | Open | Rhododendron | Understory |
| --- | --- | --- | --- | --- | --- | --- |
| Manali | 5/15/22 – 6/4/22 | 5/15/22 – 6/4/22 | 5/17/22 – 6/4/22 | 5/12/22 – 6/4/22 | 5/17/22 – 6/4/22 | 5/10/22 – 7/9/22 |
|  | n = 5 | n = 5 | n = 5 | n = 4 | n = 5 | n = 9 |
| Nain Gahar | 6/15/22 – 7/9/22 | 6/16/22 – 7/9/22 | NA | 6/15/22 – 7/8/22 | NA | 6/15/22 – 7/9/22 |
|  | n = 4 | n = 4 | n = 0 | n = 5 | n = 0 | n = 4 |

**Table S1.** Sites and sample sizes for light loggers in Fig. 6.

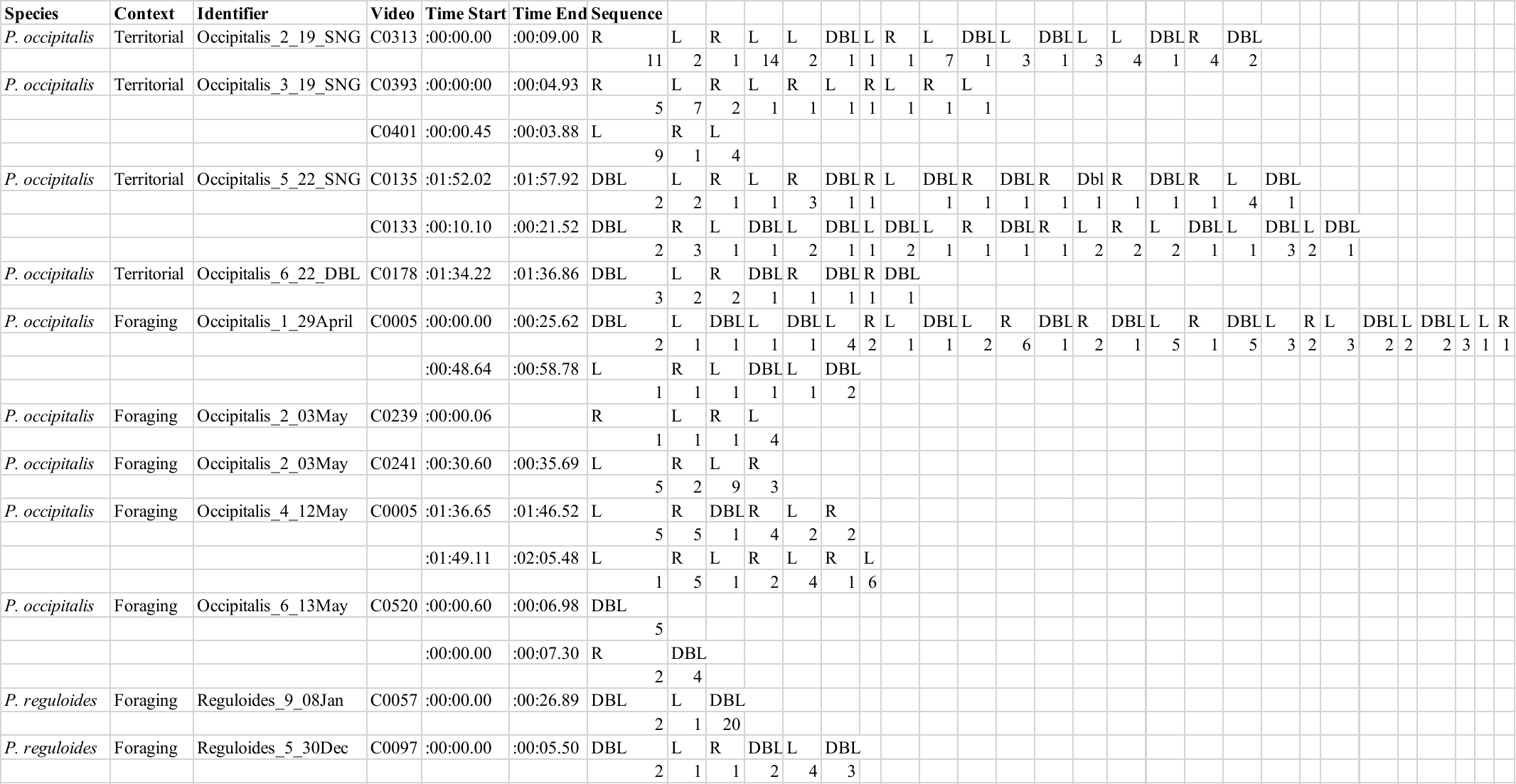
**Table S2.** The sequence of single wing flicks used by *P. occipitalis* and *P. reguloides* during territorial and foraging observations. R = right wing, L = left wing, DBL = both wings, labeled as a double wing flick.

| **Model** | **F** | **p-value** | **R^2^** | **Notes** |
| --- | --- | --- | --- | --- |
| One way ANOVA: WFS ~ Species for territorial WF | 9.04 (8,18) | **0.00*** | NA | Species differ in territorial WFS rate - however this is driven by *P. affinis* (see Tukey tests, Table S4) |
| One way ANOVA: WFS ~ Species for territorial WF without *P. affinis* | 0.44 (8,15) | 0.86 | NA |  |
| One way ANOVA: WFS ~ Species for foraging WF | 0.49 (8, 24) | 0.85 | NA |  |
| One way ANOVA: WFS ~ Context for a dataset of territorial and foraging double WF | 9.67 (1,58) | **0.00*** | NA |  |
| Linear regression Wing flick territorial ~ Wing flick foraging | 1.62 (1,7) | 0.24 | 0.19 |  |
| Linear regression Wing flick territorial ~ Wing flick foraging without *P. affinis* | 2.00 (1,6) | 0.21 | 0.25 |  |

**Table S3.** Statistical tests (ANOVA and linear regressions) for wing flicks per second (WFS), species and context (foraging vs territorial displays).

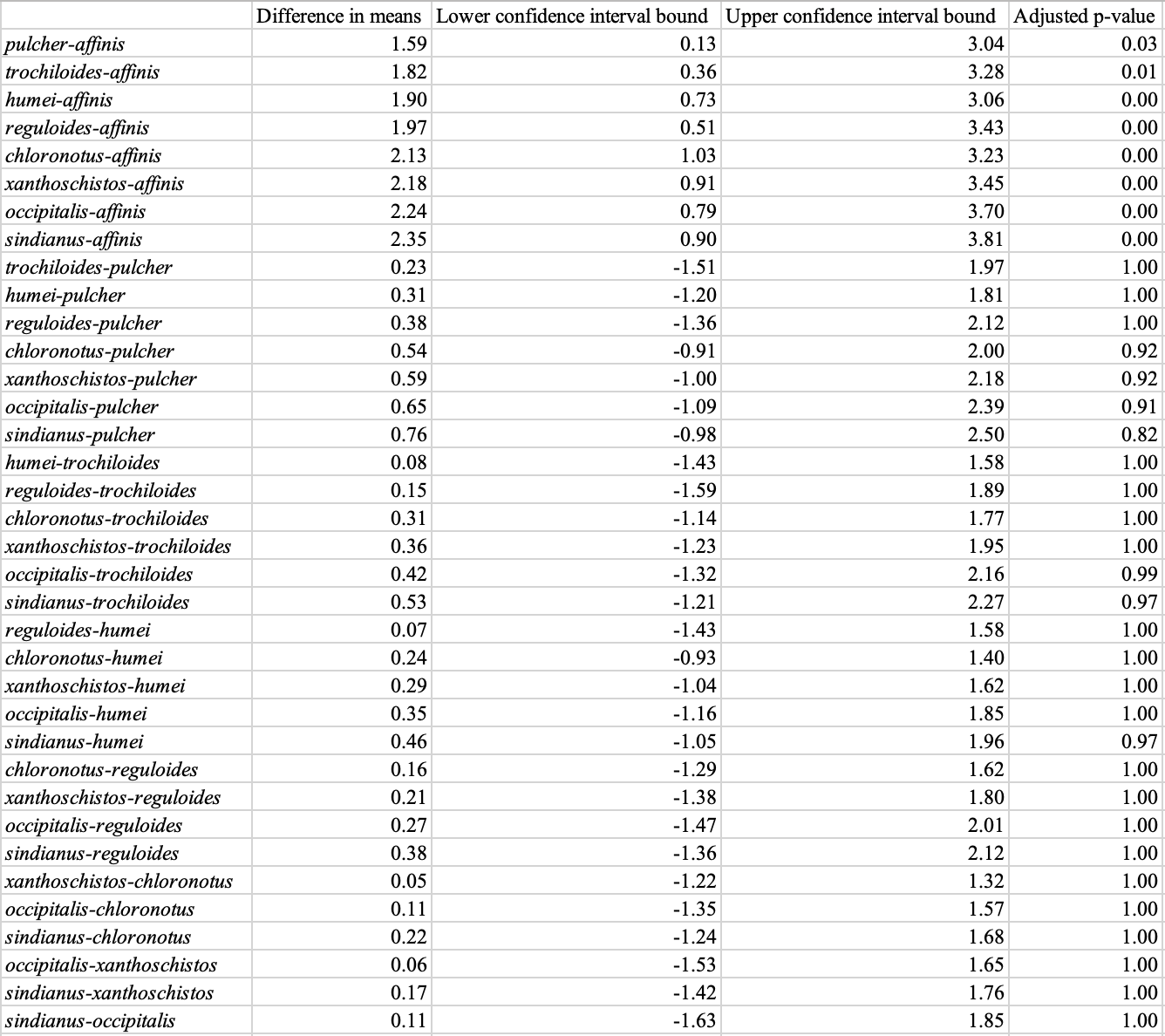

**Table S4.** Pairwise Tukey test results for species on territorial wing flick rate. All significant values include *P. affinis*, which has a wing flick rate of 0. Comparisons between all other species are nonsignificant. Only species labels are show.

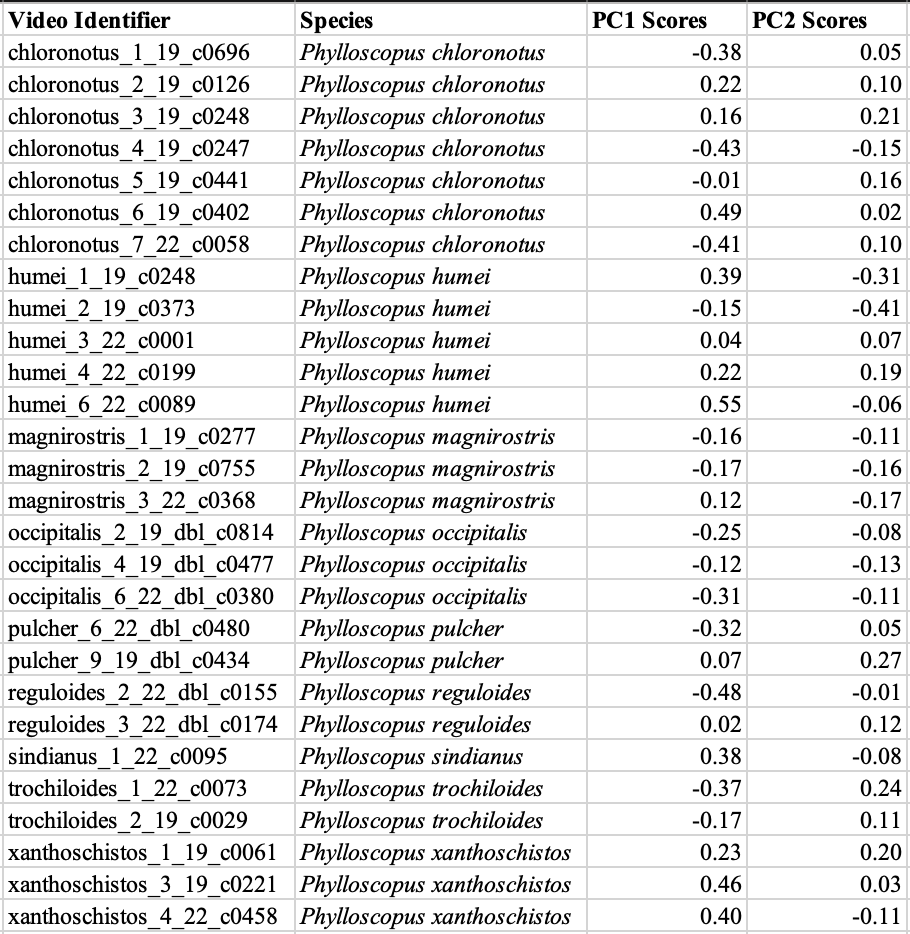

**Table S5.** PC1 and PC2 scores for double wing flick shapes

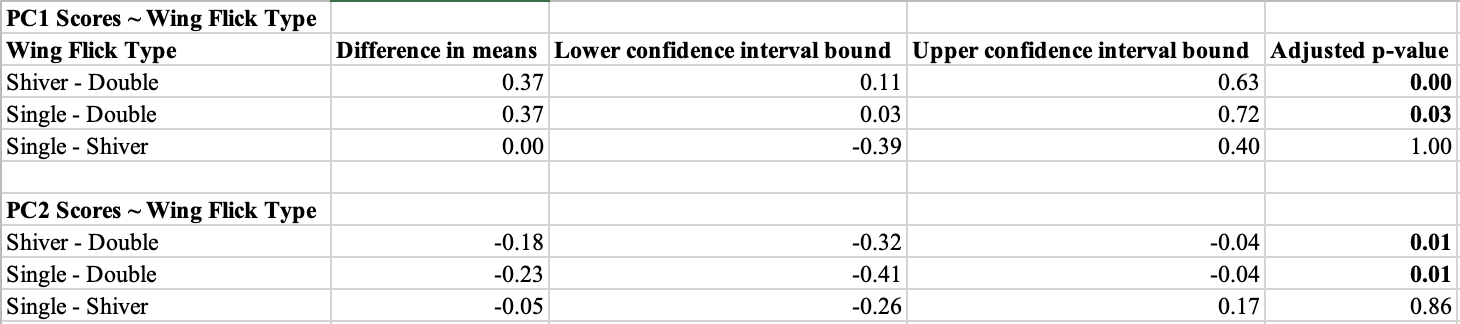

**Table S6.** Pairwise Tukey tests for PC1 and PC2 scores of wing flick shape on wing flick type. Single and shiver wing shape motions varied significantly from the double wing flick along PC1.

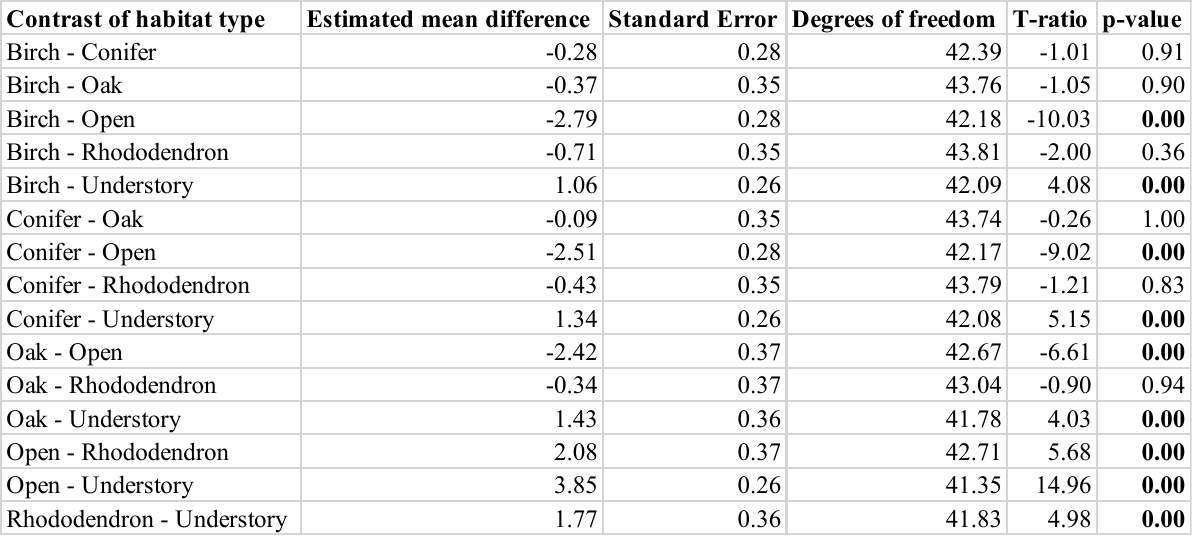

**Table S7.** Pairwise comparisons of the effects of habitat type on light (lux) from a linear mixed-effects model. Adjusted p-values were calculated using the Tukey method. Open and understory habitats are significantly brighter and darker respectively from all other habitat types.

**Supplementary Video Files:
Supp. Video 1:** Chick begging display, *P. trochiloides*, standard frame rate (60 fps).
